# Effects of assisted gene flow on the flowering onset of the annual legume *Lupinus angustifolius* L.: from phenotype to genotype

**DOI:** 10.1101/2023.01.20.524742

**Authors:** Sandra Sacristán-Bajo, Carlos Lara-Romero, Alfredo García-Fernández, Samuel Prieto-Benítez, Javier Morente-López, María Luisa Rubio Teso, Elena Torres, José María Iriondo

## Abstract

Current climate change may impede species to evolutionary adapt quickly enough to environmental changes, threatening their survival. In keystone populations, it may be necessary to consider the introduction of adaptive alleles through assisted gene flow. Considering that flowering time is a crucial trait in plant response to global warming, the objective of our study was to test the potential benefits and limitations of assisted gene flow for enhancing the evolutionary potential of *Lupinus angustifolius* L. (Fabaceae) populations through the advancement of flowering time in the context of global warming. Previous studies have shown that southern populations of *L. angustifolius* flower earlier than northern populations. We collected seeds from four populations in Spain from two different latitudes, and we established them in a common garden environment. To advance the flowering onset of northern populations, we used pollen from southern individuals to pollinate plants from northern populations, creating an F1 gene flow line. In the following season, the F1 gene flow line was self-pollinated to create an F2 self-pollination line. In parallel, individuals from the F1 gene flow line were pollinated again with pollen from northern plants, thus creating a backcross line. We also included a control line resulting from a random selection of individuals in each population in the first generation and their descendants from self-crosses in the second generation. We measured flowering onset, reproductive success and other plant traits in all individuals resulting from these lines. To characterize the effects of the assisted gene flow line at the genomic level, we carried out a gene capture analysis to sequence genes related to reproduction, growth, stress, nitrogen, and alkaloids in individuals from the F1 gene flow line and the control line in the first generation. All gene flow-derived lines flowered significantly earlier than the control line. Furthermore, plants from the F1 gene flow line produced heavier seeds and had a lower shoot growth than the control line. Genomic analyses identified 36 SNPs outliers that were associated to flowering onset, seed weight, and shoot growth. These results highlight that assisted gene flow can increase the evolutionary potential of populations by modifying the values of a specific trait. However, the modification of one trait may affect the values of other plant traits. The characteristics of the populations will have a fundamental effect on the results of assisted gene flow. Therefore, the selection of the donor population is a critical step in this process.

## Introduction

The current climate change, accelerated by human activity, is compromising the ability of organisms to survive (Langsdorf et al., 2022). Species can react to climate change through different kinds of responses, such as migration to more favorable areas, phenotypic plasticity, or evolutionary adaptation (Jump & Peñuelas, 2005; Parmesan & Yohe, 2003). Whereas migration ability can be limited for some organisms, evolutionary adaptation depends on the genetic variation, demography, and historical processes of populations (Sheth & Angert, 2016). Since, in some cases, the environment is changing faster than the rate at which species can adapt or migrate, a mismatch may occur between climate and the adaptations of organisms (Aitken & Whitlock, 2013). Therefore, it is necessary to propose strategies that can improve the adaptive potential of organisms or reduce the risk of population extinction in the context of climate change.

Assisted migration is one of the strategies that has been proposed in this line (Grady et al., 2011; Loss et al., 2011). It consists of the physical translocation of populations to habitats, outside the natural range of the species, that are expected to be more favorable for the species according to future climate predictions (Aitken & Whitlock, 2013; Vitt et al., 2010). However, assisted migration proposals have been widely debated due to the biological risks they can entail. It has been argued that this kind of intervention can cause major impacts on biotic communities, alter nutrient cycles, or disrupt ecological processes such as pollination or seed dispersal (Mack et al., 2000; Traveset & Richardson, 2006). Hybridization with other species could also occur, in addition to the possibility of the species becoming invasive or the involuntary transfer of pathogens (Loss et al., 2011; Williams & Dumroese, 2013). Furthermore, the impacts of these introductions may not be appreciable in the short term and may vary greatly over space and time (Ricciardi & Simberloff, 2009). On the other hand, gene flow can bring genetic variability to populations, allowing them to adjust to new scenarios such as climate change (Grummer et al., 2021). To facilitate gene flow, the creation of corridors has also been proposed with the purpose of improving the adaptation and conservation of organisms (Beier, 2012; Heller & Zavaleta, 2009). However, these connections are not always possible due, for example, to habitat fragmentation (Heller & Zavaleta, 2009) or natural barriers. In addition, natural gene flow is limited for some species, such as those plants that are strictly autogamous.

Assisted gene flow is a proposal that could help to solve some of these issues (Aitken & Whitlock, 2013; Prieto-Benítez et al., 2021; Wadgymar et al., 2015). It is defined as the movement of gametes or individuals between existing populations to facilitate adaptation (Aitken & Whitlock, 2013; Whiteley et al., 2015). There are some studies that have explored the possible benefits that gene flow can have on populations genetically impoverished (Morente-López et al., 2021; Prieto-Benítez et al., 2021; Sexton et al., 2011), but the possibility of using directed gene flow to improve the adaptive potential of populations in a context of climate change has been little studied. In contrast to assisted migration, assisted gene flow only implies the movement of genes or individuals between natural populations of the species already present in the ecosystem, so it is seen as a less risky strategy from an ecological point of view (Aitken & Whitlock, 2013). Another advantage of assisted gene flow versus natural gene flow resides in its wide geographical potential, as the movement of gametes can take place over very long distances. Furthermore, when done in a directional manner, the alleles that are introduced are more likely to be pre-adapted to actual or potential environmental circumstances, while natural gene flow can occur in any direction and lead to possible maladaptation (Aitken & Whitlock, 2013). However, assisted gene flow can also involve some genetic risks associated with outbreeding depression or the breakdown of sets of coadapted genes due to epistatic effects (Aitken & Whitlock, 2013; Byrne et al., 2007; Grummer et al., 2021). Outbreeding depression may also result in a loss of local adaptation, as some alleles that work well in one context may cause a fitness loss in other conditions (Frankham et al., 2011). Then, deepening our understanding of the risks and benefits of using assisted gene flow can help us to understand and improve the evolutionary capacity of populations (Frankham et al., 2017).

Within the same species, many traits of great ecological importance can vary with latitude and along environmental and temperature gradients (De Frenne et al., 2013; Milla et al., 2009). For example, phenology and climate conditions are closely related. Organisms are constantly trying to match their phenologies with the most favourable environmental circumstances (Pau et al., 2011). Phenological shifts are therefore among the most prominent impacts of climate change (Bradshaw & Holzapfel, 2009; Parmesan & Yohe, 2003), and flowering onset is a critical aspect in the adaptation of plants to climate change (Franks & Hoffmann, 2012). It has been shown that populations present differences in their flowering onset depending on the latitude, with lower latitude populations flowering earlier in most cases (Lévesque et al., 1997; Riihima et al., 2005). The fact that some of these results are obtained in common garden conditions indicates that the onset of flowering is a genetically controlled trait (Riihimäki & Savolainen, 2004) with high heritability (Franks et al., 2007). Moreover, it is also a polygenic trait in which a large number of genes are involved and has a complex regulation (Blümel et al., 2015). However, the timing of flowering onset is often correlated with other traits of great importance for plant survival and adaptation, which may constrain its evolution (Etterson & Shaw, 2001; Sacristán-Bajo et al., 2022; Walsh & Blows, 2009). Therefore, a better understanding of the genomic basis of flowering onset and the possible genetic constraints due to correlations between traits may help to design more accurately potential future assisted gene flow actions.

Given the scarce evidence available on this subject, the main objective of this study was to take an experimental approach with assisted gene flow as a way to advance the timing of flowering onset in plant populations in an effort to promote their adaptive potential to climate change, assessing the risks and benefits that it can entail. In this context, we set up an experimental study with *Lupinus angustifolius* L. (Fabaceae), using phenotypic and genomic approaches, that involved manual crosses between four populations from two climatically different areas of the Iberian Peninsula, and the characterization of the progeny in a common garden environment. In previous studies carried out with these same populations (Sacristán-Bajo et al., 2022), it has been observed that southernmost populations (which have warmer climate patterns) flower earlier than northernmost populations; therefore, we expect that the gene flow from southern populations to northern populations will produce an advance in the onset of flowering in the offspring with respect to the average individuals of the northern populations. Since the southern populations used as sources for the gene flow are genetically distinct in many traits from the recipient populations, we hypothesize that the introduction of genes originating from the south in populations from the north will also generate changes in other traits. We expect that the first generation of gene flow will produce hybrids with intermediate phenotypes. The second generation, produced by self-fertilization of the hybrids, will produce segregation of the traits, giving rise to very different phenotypes, similarly to the backcross line, obtained by the subsequent pollination of the hybrids with individuals from the northern populations. To test these hypotheses, we recorded the timing of flowering onset and several other plant traits, and compared the gene flow lines against the control line to answer the following questions: i) Is it possible to advance the flowering of northern populations through assisted gene flow from the southern populations? ii) If so, are there other traits of the individuals modified with the gene flow? iii) What is the genomic signature of the phenotypic changes brought about by the gene flow lines?

## Materials and Methods

### 1. Study species and source populations

The blue lupine (*Lupinus angustifolius* L.) is an annual legume distributed throughout the Mediterranean basin, which has been domesticated as a crop and is grown in many places around the world (Castroviejo & Pascual, 1993). This plant can reach up to more than 100 cm in height and has characteristic palmate leaves divided in 5-9 leaflets. Its flowers are hermaphrodite, with an inflorescence that can have up to 30 flowers. The fruit is a dehiscent legume with 3-7 seeds (Clements et al., 2005). The species mainly self-pollinates before its petals open (Wolko et al., 2011), with outcrossing estimates below 2 % (Dracup & Thomson, 2000). Depending on latitude and environmental conditions, flowering occurs between March and August (Castroviejo, 1999), since its flowering onset is influenced by photoperiod and temperature (Rahman & Gladstones, 1974).

For our study, we selected four populations distributed by pairs in two regions of contrasting climatic conditions, the northern one located in Salamanca (Central Spain), and the southern one located in Badajoz (South Spain) (Figure 1, Table S1). The distance between the two regions is around 300 km, and between the populations within each region is less than 20 km. Both regions have similar annual precipitation, but the southern one has higher mean, minimum and maximum temperatures and, consequently, experiences higher water deficits. We collected seeds in each population from at least 98 genotypes (mother plants) located at least one meter apart from each other.

**Figure 1.**
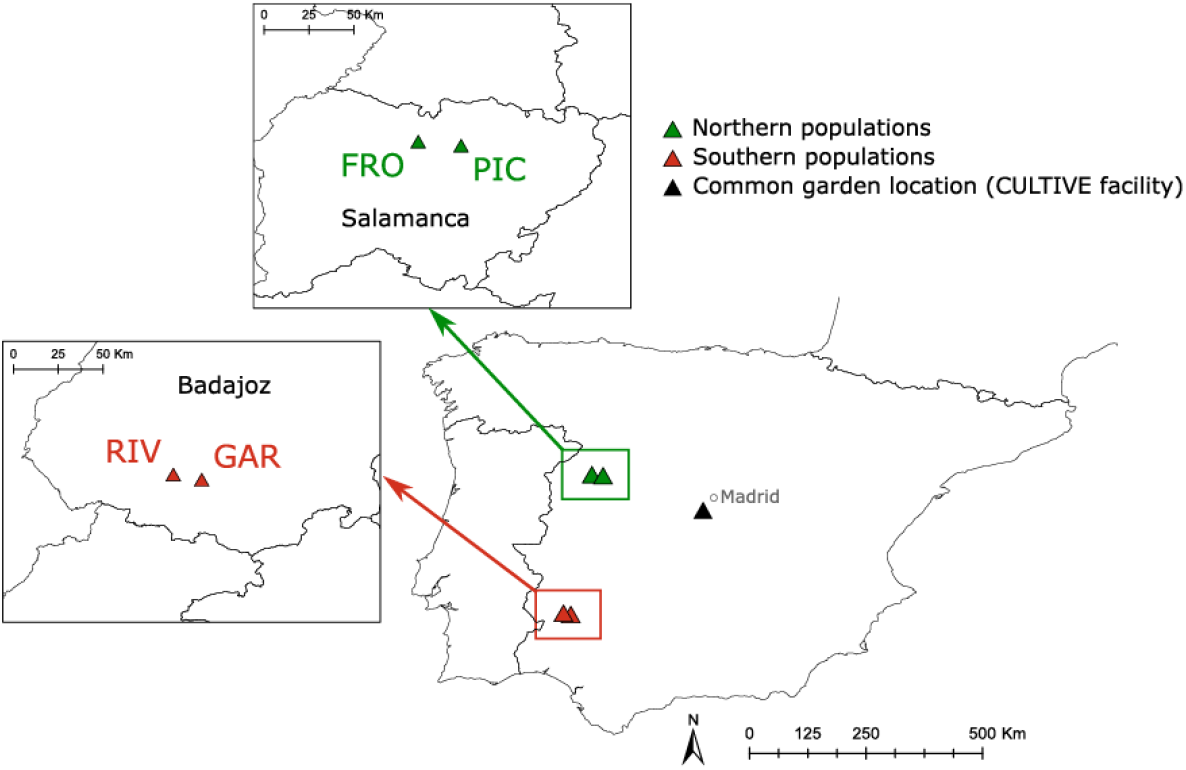
Location of the populations of *Lupinus angustifolius* L. and of the common garden used in the experiment.

### 2. Common garden and gene flow experiment

The experiment was carried out in a greenhouse of the CULTIVE facility (https://urjc-cultive.webnode.es/) at Rey Juan Carlos University (Móstoles, Madrid). Inside the greenhouse, the temperature varied between 1 to 25 degrees Celsius, and plants only received natural light. In November 2016, three seeds from each maternal genotype were scarified to ensure germination and sown in a 6 L pot, following the same protocol described in Sacristán-Bajo et al. 2022. In spring 2017, the pots were transferred outside of the greenhouse to the CULTIVE experimental field and were distributed in a randomized block design where the plants from the different populations were evenly represented in each block. The substrate in the pots was kept at field capacity with a system of drip irrigation. Before the plants flowered, four siblings from 25 % of random genotypes per population were selected and their inflorescences in their main shoot were bagged to obtain seeds derived from self-pollination. This first growing season was only used to eliminate maternal effects.

In November 2017, seeds separately collected from each individual were sown in the same way and with the same conditions as described above, and the resulting plants were transferred to the CULTIVE experimental field in February 2018. During the flowering season, manual between-population crosses were carried out to create an ‘F1 gene flow line’ (hereafter, GFL). Plants from the northern region were pollinated using pollen from plants from the southern region, matching the RIV population with the PIC population and the GAR population with the FRO population. All possible crosses between these two pairs of populations (considered as replicates) were performed considering the need of having overlapped flowering periods between their individuals. The procedure to carry out manual crosses was the same as that described in Sacristán-Bajo et al. (2022), based on the emasculation of individuals of the northern region and their subsequent pollination with pollen from individuals from the southern region. Seeds produced were separately collected for each mother plant. In parallel, for each population, the 25 % of the genotypes were naturally self-crossed, and their seeds were separately collected to generate the ‘control line’ (hereafter, CFL). In the 2018-2019 season, seeds were sown, and seedlings were cultured and transferred outdoors in the same way as described above, containing, for each population, individuals from the CFL (64 individuals for FRO population and 65 individuals for PIC population) and GFL (33 individuals for FRO population and 43 individuals for PIC population). In the flowering season of 2019, the individuals of the GFL were manually pollinated using individuals of the CFL of the northern populations as pollen donors, creating a ‘backcross line’ (hereafter, BCL). Additionally, an ‘F2 self-pollination line’ (hereafter, SPL) from the GFL was generated by self-pollination. The seeds of these lines, as well as those derived from the self-pollination of CFL, were again separately collected for each mother plant. In the 2019-2020 season, the seeds from these lines were sown and the resulting seedlings were grown and transferred outdoors as indicated above. To summarize, a diagram of the complete process is shown in Figure 2. The manual crossings between individuals from paired northern-southern populations were constricted by the need to have overlapping flowering periods between pairs of individuals. The success of the manual crossings performed was low (around 7 %). As a result, 21 mother plants from both populations were successfully crossed to obtain eight different genotypic crosses in FRO population and 13 different genotypic crosses in PIC population.

**Figure 2.**
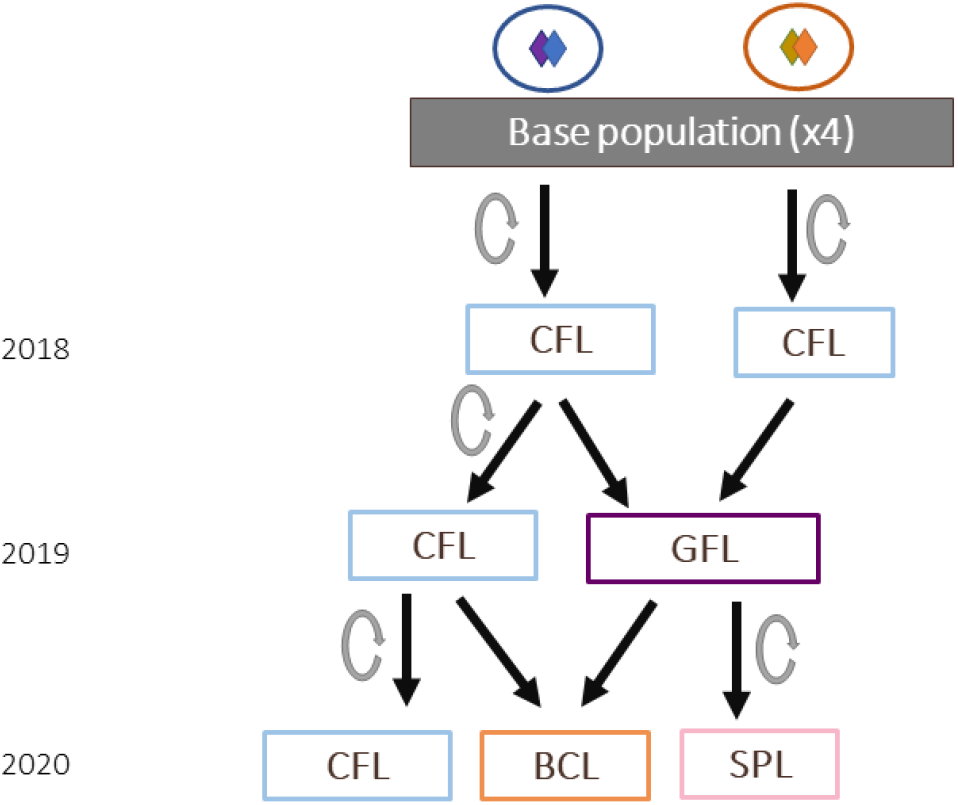
Flowchart describing the different lines performed with individuals of four populations of *Lupinus angustifolius* through time. Blue and orange diamonds represent the populations of the northern and southern regions, respectively. CFL: control line; GFL: F1 gene flow line; BCL: F2 backcross line; SPL: F2 self-pollination line. Gray circled arrows indicate that the individuals of that line were self-crossed. Black arrows denote the transmission of gametes for the next generation. The years indicated on the left correspond to the flowering season in which adult plants of the indicated lines had been grown and the crosses performed in that season are represented below with the arrows.

### 3. Traits measurement

The day of flowering initiation was recorded for each plant as the day when the first purple flower of the main inflorescence was clearly visible. Thus, we defined the flowering onset variable as the number of days between the date of sowing and the flowering start date. We also calculated the height of the plants (cm) at the flowering peak (when most plants had bloomed) by measuring the distance from the soil level to the base of the main inflorescence, and also, the length of the main inflorescence. We estimated the number of fruits per plant based on the total number of floral scars at the end of the season. The average number of seeds per fruit was determined by counting the seeds in 15 different fruits per plant. The number of seeds per plant was calculated by multiplying the number of fruits per plant by the average number of seeds per fruit. The individual weights of 10 random seeds from each plant were used to calculate the mean seed weight. The mean seed weight and the number of seeds per plant were used as proxies for determining plant fitness.

The central leaflet from eight fully developed leaves belonging to the lateral branches was gathered to determine the specific leaflet area (SLA) and dry matter content (LDMC). The fresh leaflets were weighed immediately in a Kern ABJ 120-4M analytical balance (Kern & Sohn GmbH, Albstadt, Germany). The leaflets were then placed in water-soaked filter paper and stored in plastic bags before being refrigerated overnight at 4 °C. We weighed the leaflets again the next day to get the turgid weight and used a foliar scanner Li-3000C (Li-Cor, NE, United States) to measure the area of the leaflets. Finally, the leaflets were dried for at least 72 hours in a 60 °C oven before being weighed again to determine their dry weight. SLA was calculated by dividing the area of a leaflet by its dry weight (Rosbakh et al., 2015). LDMC was determined by dividing the leaflets dry weight by its saturated weight (Wilson et al., 1999). At the start of flowering and at the end of the culture cycle, we measured the length of the plant from its base to the first flower. The difference between these two values was used to estimate the shoot growth of each individual. We also measured the aboveground biomass of each plant at the end of the culture cycle.

The flowering onset was measured for the years 2019 and 2020, but due to the pandemic lockdown, the rest of the traits were only measured for the year 2019.

### 4. Phenotypic analyses

We used the R statistical environment version 4.1.1 to conduct all the statistical analyses mentioned below (http://r-project.org). We applied linear and generalized linear mixed models (hereafter, LMMs and GLMMs) to analyze the effect of the F1 gene flow line as well as the F2 self-pollination line and the backcross line on flowering onset and the rest of the traits. Therefore, for each trait, we included the *line* (CFL and GFL for the year 2019, and CFL, SPL and BCL for the year 2020) and the *population* (FRO and PIC) as fixed effects, and *genotype* (mother plant) as random effect. For the flowering onset variable, we used a Poisson error distribution, and for the rest of the variables, we used a Gaussian error distribution. Diagnostic plots were used to visually inspect the model residuals for normality and variance homogeneity. We tested the interaction between the variables *line* and *population*. As the interaction was not significant, we decided not to include it in the models. The *glmer* and *lmer* functions from the lme4 package version 1.1-27.1 were used to fit the GLMMs and LMMs (Bates et al., 2015). The *Anova* function from the car package version 3.0-11 was used to determine the significance of each fixed effect (Fox & Weisberg, 2011). If necessary (as for the flowering onset in 2020), Tukey *post hoc* analysis from the *emmeans* function from the emmeans package version 1.6.3 was used to calculate differences between lines (Lenth, 2019). R^2^ values were calculated using the *summ* function from the jtools package version 2.2.0 (Long, 2019). The *corrplot* function version 0.90 from the corrplot package was used to plot correlations between flowering onset and the other traits (Wei et al., 2017).

### 5. Genomic analyses

#### DNA extraction and selection of candidate genes

Leaf material was collected for DNA extraction in 2019. Therefore, it corresponded to the individuals of the CFL and GFL lines that were phenotyped. Leaves from a total of 60 individuals were collected, 30 of each line (CFL and GFL), and 15 of each population (FRO and PIC) within each line. We used DNeasy Plant minikit (QIAGEN, Valencia, USA) to extract and isolate DNA.

We designed a gene capture experiment, taking the annotated *Lupinus angustifolius* genome from the National Center for Biotechnology Information (NCBI) as starting point (GenBank accession: PRJNA398717). This genome contains all coding sequences (CDs) belonging to the *L. angustifolius* genome. FullLengtherNext software (Lara et al., 2007) and the *L. angustifolius* genome were used to perform a Blast analysis and obtain the biological function (i.e., gene ontology terms) for each sequence in *L. angustifolius*. We selected 73 gene ontology terms related to reproduction, growth, stress, nitrogen, and alkaloids. After that, we used Go.db package in R software to create a list with all the gene ontology terms parent and children of these gene ontology terms. Finally, we filtered the file which contains all coding sequences to obtain the candidate genes of interest, based on the list of gene ontology terms. These sequences were used as probes to carry out the targeted sequencing.

#### Sequencing and Single Nucleotide Polymorphism (SNP) calling

The extracted DNA was sent to IGATech (Udine, Italy). The quality of the genomic DNA was checked using Qubit 2.0 Fluorometer (Invitrogen, Carlsbad, CA) and NanoDrop 1000 Spectrophotometer (Thermo Fisher Scientific, Waltham, Massachusetts). Libraries for target enrichment of ∼3 Mb of *Lupinus angustifolius* genomic material were produced using the Roche Sequencing Solutions’ ‘SeqCap EZ – HyperPlus’ kit (Roche Sequencing Solutions, Pleasanton, CA) with 200 ng/L of input DNA.

After that, base calling and demultiplexing was carried out with Illumina bcl2fastq v2.20. For the following steps, the software used were ERNE v1.4.6 (Del Fabbro et al., 2013) and Cutadapt (Martin, 2011) for quality and adapter trimming, BWA-MEM v0.7.17 (Li & Durbin, 2009) for the alignment to the reference genome, and Picard tools (http://broadinstitute.github.io/picard/) to produce on-target alignment statistics and metrics.

SNP calling was performed on the entire sample simultaneously with gatk-4.0 (Depristo et al., 2011). This step allowed the initial identification of ca. 41,419 SNPs. Raw SNP data were filtered using VCFtools v0.1.14 (Danecek et al., 2011), and the *vcffilter* function of VCFLIB (Garrison et al., 2021). Only biallelic SNPs with fewer than 10 % missing data were kept. Indels were also removed from the dataset. SNPs were then filtered following the hard filtering suggested by GATK’s user guide (https://gatk.broadinstitute.org/). Hence, SNPs were filtered based on their quality depth (QD > 2), Phred scaled P-value using Fisher’s Exact Test to detect strand bias (FS < 60), Symmetric Odds Ratio of 2×2 contingency table to detect strand bias (SOR < 3), Square root of the average of the squares of the mapping qualities (MQ > 40), Z-score from Wilcoxon rank sum test of Alt vs. Ref read mapping qualities (MQRankSum > -12.5), u-based z-approximation from the Rank Sum Test for site position within reads (ReadPosRankSum > - 8) and depth coverage (DP >10). This stringent filtering reduced the SNP dataset to 34,026 SNPs. Finally, SNPs in high linkage disequilibrium were filtered using r^2^ of 0.6 as the cut-off point, which generated a final dataset of 22,802 SNPs.

#### Detecting signatures of selection

We applied a sequential strategy to identify highly divergent loci between the CFL and the GFL lines. We first calculated allele frequency differences (AFDs) between the CFL and the GFL at the individual SNP level and selected those SNPs that had experienced an allele frequency change in the same direction in both CFL vs. GFL comparisons. We then selected those SNPs with significant AFDs by applying a Fishers’s exact test (Fisher, 1970). Secondly, pairwise F_ST_ values (CFL vs. GFL) were calculated for each SNP. Statistical significance of F_ST_ values was tested for each locus by the chi-square test, *x*^*2*^*= 2NF*_*ST*_*(k - 1)*, with *(k - 1)(s - 1)* degrees of freedom, where *N* is the total sample size, *k* is the number of alleles per locus, and *s* is the number of populations (Workman & Niswander’, 1970). We only considered that an SNP showed divergent patterns of differentiation when it was selected as an outlier by both *F*_ST_ analyses and at the same time it showed consistent AFDs in the two pairs of CFL vs. GFL comparisons. Lastly, these highly divergent loci underwent an individual genotype-phenotype validation (Chen et al., 2022). For this purpose, a linear mixed model with random family effects was fitted using the traits flowering onset, seed weight, and shoot growth as dependent variables, the genotype of each SNP as a three-level explanatory factor (homozygous for the minor allele, homozygous for the major allele and heterozygous), individual as a random factor and a kinship matrix as a random genetic effect to control for kinship effects. This validation allowed us to detect those SNPs with a large effect on the phenotype. F_ST_ values and allele frequencies were calculated using VCFtools v0.1.14. Kinship matrix was calculated using the centered-IBS method implemented in TASSEL v5.2.81 (Bradbury et al., 2007). Linear mixed models were fitted using the *lmekin* function implemented in *coxme* R package (Therneau, 2020).

## Results

### Flowering onset

Significant differences in flowering onset were found between the GFL and the CFL in 2019 and also between the BCL and SPL and the CFL in 2020 for both populations (Figure 3). In 2019, plants from the GFL flowered an average of 7 days earlier than control plants in FRO population, and an average of 8 days earlier in PIC population (*X*^2^ = 20.21, p < 0.001, Df = 1) (Figure 3a, Supplementary material Table S2, S3 and S4). In 2020 the SPL flowered 6 days earlier than the CFL in both populations, and the BCL flowered 12 days earlier than the CFL in FRO population (*X*^2^ = 13.00, p = 0.001, Df = 2) (Figure 3b, Supplementary material Table S2 to S4). The values of R^2^ indicated that the fixed effects explained 10 % of the variation, and the random effects explained 2.2 % of the variation in the year 2019. In the year 2020, fixed effects explained 25.3 % of the variation, whereas the random effects explained 16.4 % (Supplementary material Table S3). Posterior mean values, standard errors, and 95 % confidence intervals for each line are shown in Supplementary material Table S5

**Figure 3.**
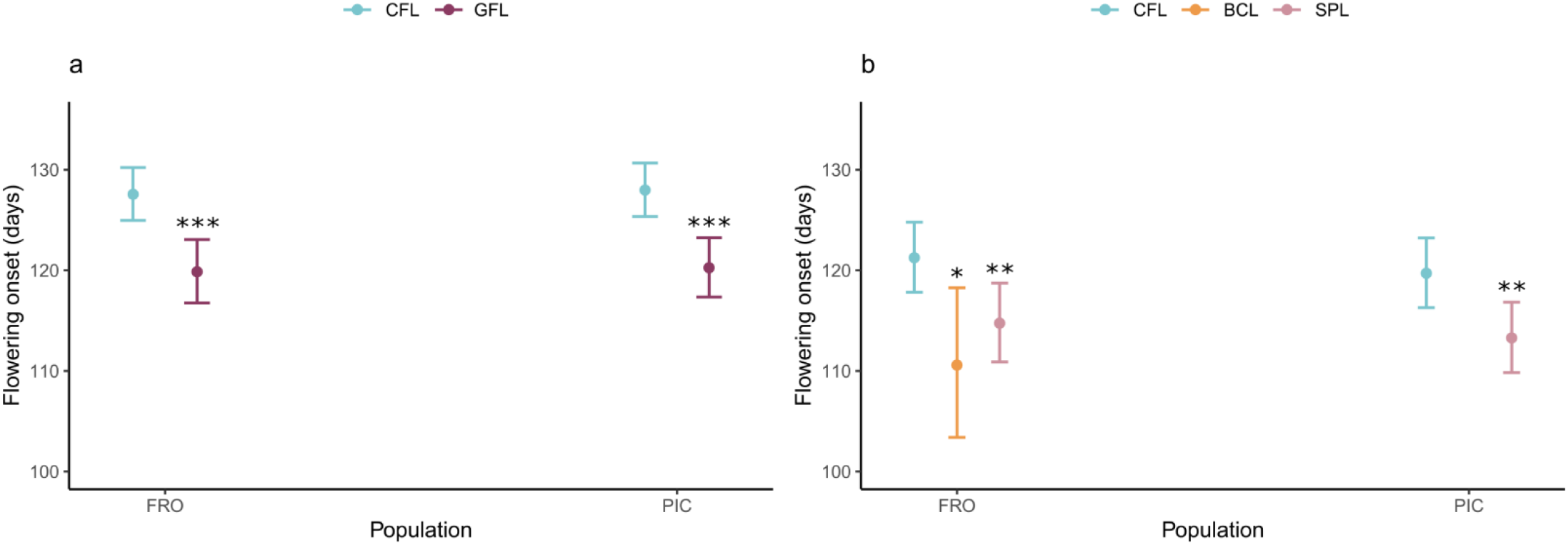
Effect of the different lines on advancing flowering onset of *Lupinus angustifolius*. a) effect of the gene flow line (2019), b) effect of the F1 self-pollination line and backcross line (2020). CFL: control line; GFL: F1 gene flow line; BCL: F2 backcross line; SPL: F2 self-pollination line. The values for flowering onset correspond to the estimates obtained from the GLMM model. Dots and bars represent the predicted mean from the GLMM model with a Poisson distribution and 95 % confidence intervals. Significant differences (p < 0.05) determined by Tukey test between the created lines and the control line are marked with asterisks (*p < 0.05; **p < 0.01; ***p < 0.001).

### Reproductive success

No significant differences were found between the GFL and the CFL for seed number (*X*^2^ = 1.38, p = 0.24, Df = 1) (Figure S1a, Supplementary material Table S4), but significant differences were obtained for seed weight (*X*^2^ = 29.96, p < 0.001, Df = 1). Seeds from GFL were heavier than those from CFL (Figure 4a, Supplementary material Table S2, S3 and S4). In addition, significant differences were found between the two populations both for seed number (*X*^2^ = 4.64, p = 0.03, Df = 1) (Figure S1a, Supplementary material Table S4) and seed weight (*X*^2^ = 8.20, p = 0.004 Df = 1) (Figure 4a, Supplementary material Table S4). For seed number, fixed effects explained 12.5 % of the variation, whereas the random effects explained 8.6 %. For the seed weight, fixed effects explained 38.8 % of the variation, and the random effects explained 16.3 % (Supplementary material Table S3). Posterior mean values, standard errors, and 95 % confidence intervals for each line are shown in Supplementary material Table S5.

**Figure 4.**
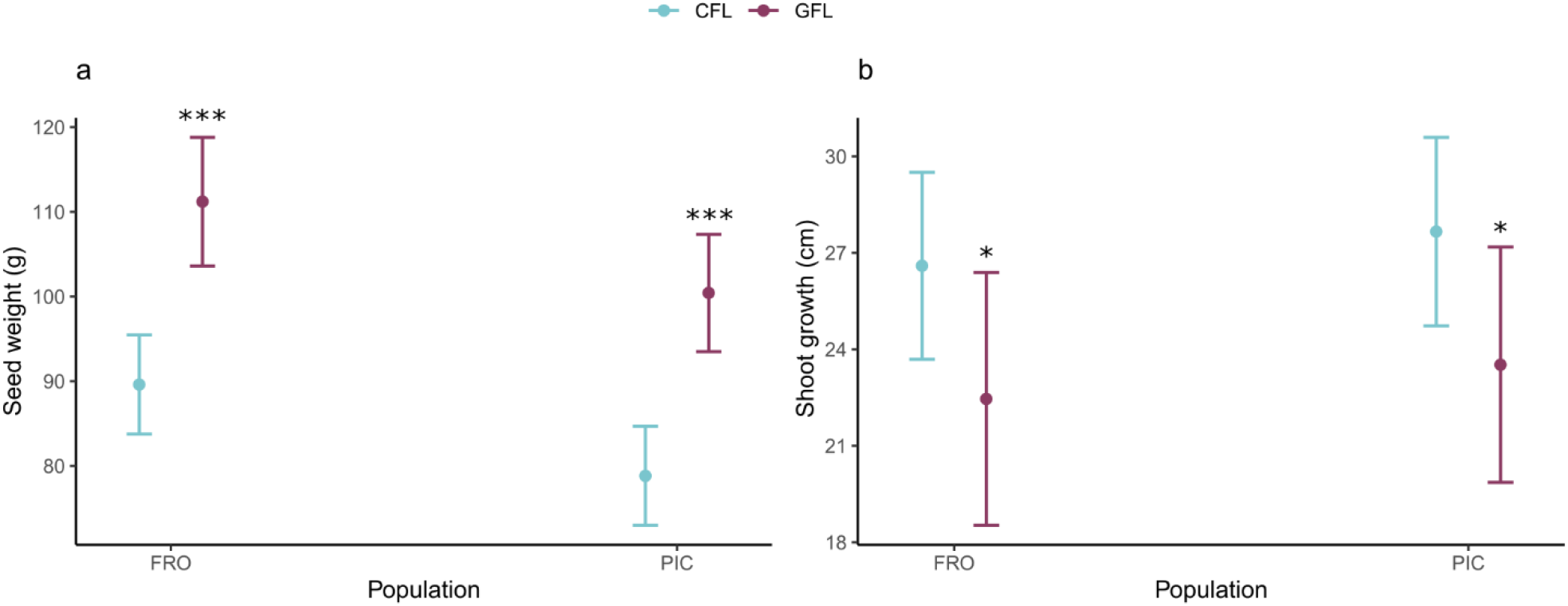
Effect of the gene flow line on the seed weight and shoot growth of *Lupinus angustifolius*. a) effect on the seed weight, b) effect on the shoot growth. CFL: control line; GFL: F1 gene flow line. Dots and bars represent the predicted mean from the LMM model with a Gaussian distribution and 95 % confidence intervals. Significant differences (p < 0.05) determined by Tukey test between the gene flow line and the control line are marked with asterisks (*p < 0.05; **p < 0.01; ***p < 0.001).

### Other plant traits

Regarding other plant traits not related to the onset of flowering or reproduction, significant differences were only found between the GFL and the CFL for shoot growth (*X*^2^ = 4.11, p = 0.04, Df = 1). Plants from the GFL had lower shoot growth than those from the CFL (Figure 4b, Supplementary material Table S2, S3 and S4). Depending on the studied traits, R^2^ values indicated that fixed effects explained between 0.3 % and 6.1 % of the variation, and random effects, between 0 % and 46.8 % (Supplementary material Table S3). Posterior mean values, standard errors, and 95 % confidence intervals for each line are shown in Supplementary material Table S5.

Different correlations were found between flowering onset and other plant traits for CFL and year 2019 (Figure 5). For the FRO population, plants that flowered earlier showed an increase in their height, biomass, and the weight of seeds they produced, and a decrease in their shoot growth (Figures 5a). For the PIC population, plants that flowered earlier showed an increase in seed number and weight, and a reduction in height and shoot growth (Figure 5b).

**Figure 5:**
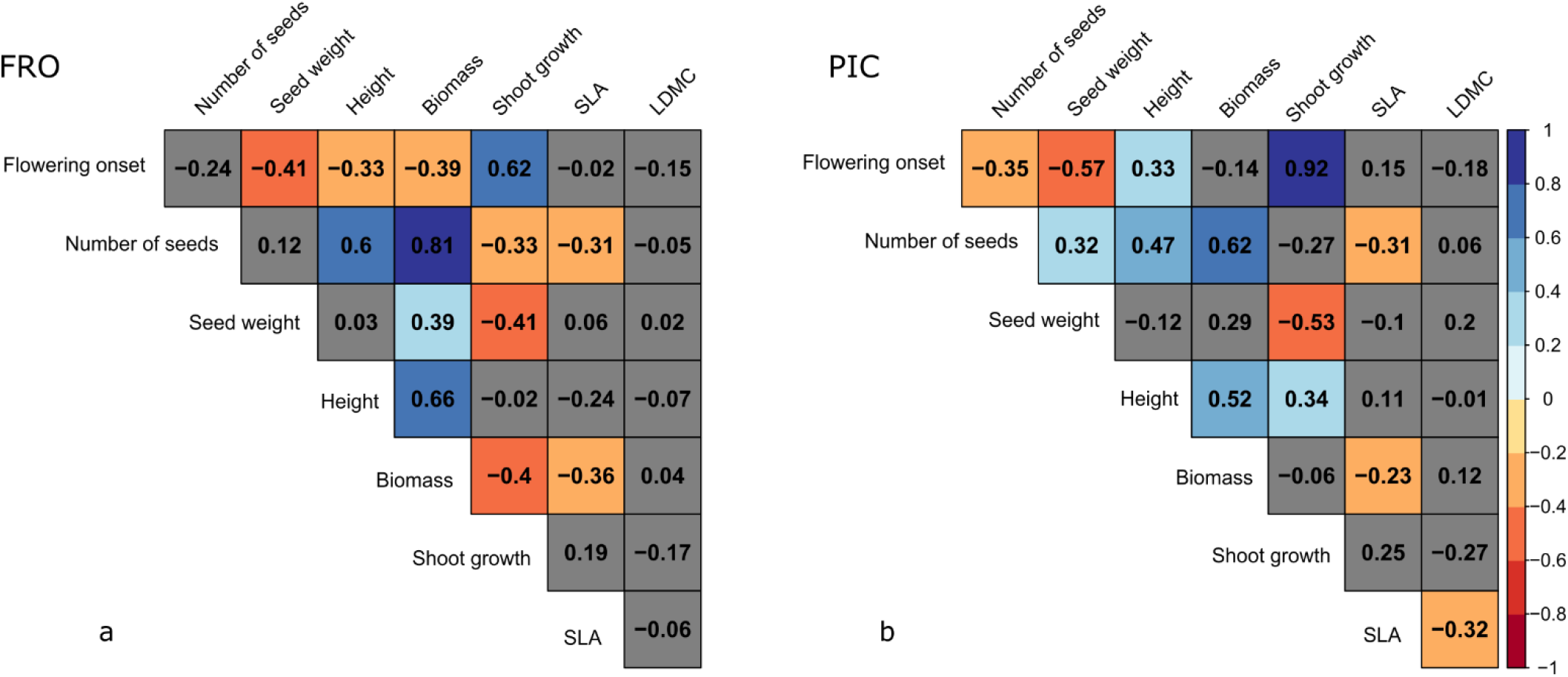
Correlations between flowering onset and other plant traits for control line and year 2019. a) correlations for FRO population. b) correlations for PIC population. Positive correlations are represented in cold colors, while negative correlations are represented in warm colors. Non-significant correlations (p > 0.05) are represented in gray.

### Loci under selection

We identified 36 SNPs that revealed divergent patterns of differentiation because they were identified as outliers both by F_ST_ analyses and exhibited consistent AFDs in the two groups of CFL versus GFL comparisons (Supplementary material Table S6). In addition, these 36 SNPs had a significant effect on flowering onset, seed weight, and shoot growth (Supplementary material Figures S2, S3 and S4) and a great change in the allele frequencies is observed between the CFL and the GFL (Supplementary material Figure S5). These SNPs were distributed on 11 of the 20 chromosomes of *Lupinus angustifolius* (Figure 6). The functional annotation revealed that the loci to which SNPs identified as outliers are related to different biological processes. Six of them were related to reproduction, another six of them were related to growth, nine of them were related to abiotic stresses, and 13 of them related to flowering (See Supplementary material Table S7).

**Figure 6:**
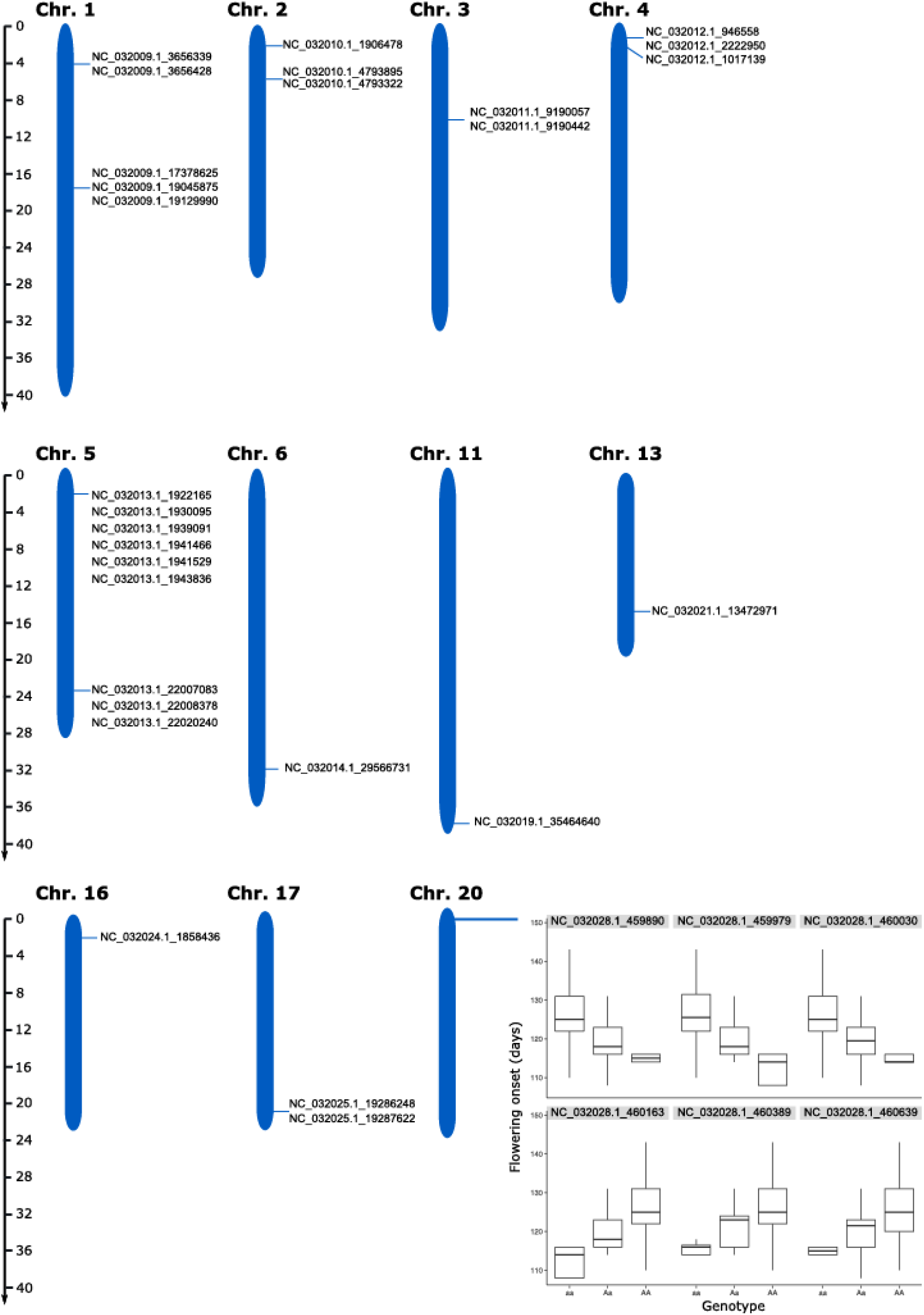
Localization of the SNPs identified to be under selection and with a significant effect on flowering initiation, seed weight, and shoot growth on the chromosomes of *Lupinus angustifolius* and detail of the SNPs localized in chromosome 20 as an example. The name of the NCBI Reference Sequence and the position of the SNP in number of base pairs are indicated in red and separated by a low bar.

## Discussion

The different lines derived from assisted gene flow produced a significant flowering advance in both populations of *Lupinus angustifolius*. As expected, the assisted gene flow also caused modifications in other plant traits, such as seed weight and shoot growth. The genomic analysis identified 36 SNPs with contrasting frequency differences between the GFL and CFL in both populations. We found that these 36 SNPs significantly explained variation in flowering onset, seed weight and shoot growth, supporting the genetic basis of flowering advancement and the presence of genetic correlations between flowering onset, seed weight and shoot growth detected at the phenotypic level. The present observations highlight the importance of evaluating the effects of gene flow from an integrative perspective.

### Effects of artificial gene flow on plant phenology, reproductive success, and functional traits

Sacristán-Bajo et al. (2022) observed that southern populations flower earlier than northern populations when grown together in a common garden. Therefore, it is reasonable to expect that hybrids from artificial crosses of northern mother plants with pollen from southern populations (GFL) will flower earlier than their respective northern population controls. It has already been suggested that natural gene flow may influence genetic changes related to climate change in plants (van Dijk & Hautekèete, 2014). These authors compared changes in flowering onset in a common garden environment in populations of *Beta vulgaris* from different latitudes and two collection years (20 years apart). They found evidence of genetic changes in response to climate change, as lower latitude populations delayed flowering, while higher latitude populations advanced flowering. Similarly to our observations, (Bontrager & Angert, 2019) found that gene flow from historically warmer populations enhanced the adaptive responses of colder populations in a context of rising temperatures. Prieto-Benítez et al. (2021) also conducted artificial gene flow experiments with the objective of modifying flowering onset in *Silene ciliata* Pourr. (Caryophyllaceae), a perennial mountain plant. Contrary to our results, they observed that gene flow from populations that flowered earlier did not advance the onset of flowering in the recipient populations, but rather delayed it. This shows that the effects of assisted gene flow on flowering onset can be complex and species or context dependent. Outbreeding depression or epistatic effects between genes (Blümel et al., 2015; He et al., 2019; Prieto-Benítez et al., 2021) could make gene flow difficult to predict or ineffective. This could cause the backcross line (BCL) to produce a higher flowering advance than the F1 gene flow line, as the adverse effects that gene flow can cause in the first generation could be mitigated with the addition of greater genome representation from the original populations (northern populations). The possibility of carrying out successive generations of backcrossing with the population of origin while selecting for the progeny with early flowering should be considered, followed by the reintroduction of these individuals into their populations of origin.

Gene flow may not only affect the trait of interest to be shifted (in this case, flowering onset), but this modification may lead to changes in other traits, including traits related to reproductive success (Aitken & Whitlock, 2013; Morente-López et al., 2021; Prieto-Benítez et al., 2021). Considering that the southern populations are known to produce fewer but heavier seeds than the northern populations (Matesanz et al., 2020; Sacristán-Bajo et al., 2022), the GFL produced heavier seeds than the CFL individuals, making the hybrids more similar to the individuals of the southern populations. In our study, the GFL showed lower shoot growth and also a tendency to lower SLA than the CFL. Correlation analyses between the studied traits also support the existence of these same associations between flowering onset, seed weight and shoot growth. For both populations, flowering onset correlated in the same direction for these same traits (negatively with seed weight and positively with shoot growth). This indicates again that early flowering plants have higher seed weight and lower shoot growth. In addition, these changes have also been observed at the genomic level (see next section). Several other studies have shown that gene flow leads to changes in different plant traits. For example, Chacón-Sánchez et al., (2021) reviewed the effects of gene flow between cultivated and wild types of several species of the genus *Phaseolus* (Leguminosae). Morphological, seed and other traits resulted to be influenced by gene flow events and with important consequences for the species performance. In a context of climate change, shifts towards traits more similar to plants from the southern areas may be an adaptive advantage. In the same direction as our findings, Matesanz et al. (2020) observed that populations in southern areas and those subjected to drought treatment had higher seed weight, lower growth rate and thicker leaves. Plant resources are limited, and when destined for one purpose, they cannot be used for others (Reich, 2014). Therefore, the production of heavier seeds could ensure their survival in more unfavorable environments, such as the southern sites with drier conditions (Leishman et al., 2009; Metz et al., 2010), whereas lower shoot growth and lower SLA could indicate a more efficient investment of resources (Wrightet al., 1994). These results obtained in our study and in other studies reinforce the idea that the modification of certain target traits through gene flow will also come associated to changes in other traits, because biological characteristics of organisms are intricately linked, and they must be interpreted as a whole (Sobral, 2021).

### Genomic effects of the artificial gene flow

The combination of genome-wide studies with phenotypic characterization is essential to identify regions related to adaptive variation (Evans et al., 2014). In our study, we identified 36 highly divergent SNPs between control and gene flow line, both by F_ST_ and AFDs analyses, indicating that these SNPs have undergone a process of change at the genomic level due to assisted gene flow. Furthermore, we determined that flowering onset, shoot growth and seed weight are partially explained by these same SNPs. This reinforces the idea that assisted gene flow has had an impact on these traits at the genetic level, already observed in the phenotypic study carried out in the common garden experiment.

Some of these 36 loci have also been identified in other studies and have been related to growth, flowering, or abiotic stresses. For example, the axial regulator YABBY 1-like has been related to the flower development (Kumaran et al., 2002; Siegfried et al., 1999), similarly to our findings. We have related the xyloglucan endotransglucosylase/hydrolase protein to the stamen development, whereas other authors have related this protein to root growth (Maris et al., 2020) or abiotic stresses such as drought, salinity or cold temperatures (Keun et al., 2006). Nazari et al., (2020) exposed wheat (*Triticum aestivum*) to drought conditions, and using proteomics, related 3-oxoacyl protein to water stress. In our case, we found this protein related to cold responses. The protein Flowering Locus T (FT) has been identified in *Arabidopsis* and other species, including legumes, as one of the main components that promotes the onset of flowering (e. g. Kardailsky et al., 1999; Pin & Nilsson, 2012; Weller & Ortega, 2015). As in our case, this protein is known to play a role in different pathways, such as photoperiod. Related to this, the protein EBS regulates chromatin expression, controlling the expression of genes such as FT (López-González et al., 2014). It is also involved in other processes, such as organ development or seed dormancy (Gómez-Mena et al., 2001; Piñeiro et al., 2003). The allele frequency of the FT gene increased in the gene flow line compared to the control line, while SNPs related to the EBS protein decreased in frequency in the gene flow line (Supplementary material Figure S5). Thus, we found that changes produced at the phenotypic level through gene flow (such as earlier flowering) are also explained at the genomic level.

This is one of the first studies that has combined the evaluation of the use of assisted gene flow with a genomic approach. Our findings demonstrate that including genomic analyses in assisted gene flow studies provides much more accurate information about changes occurring at the genome level. In addition, the identification of these genes opens the door to the development of specific markers to identify early flowering genes in the species that may also allow the production of heavier seeds.

### Final conclusions

Assisted gene flow through human actions can contribute to the adaptations of populations to climate change by providing genetic variation (Grummer et al., 2021). With our experiments, we have observed that assisted gene flow has made possible to alter characteristics that are crucial for the adaptation of species to climate change, such as flowering onset, which could imply an enhancement of its adaptive potential in this situation. In our case, the modification of flowering onset was also associated with higher seed weight produced by plants in the F1 gene flow line and lower shoot growth. In addition, genomic analyses confirmed that these changes in phenotype due to gene flow are also observed at the genome level. Although this could be interpreted as a better adaptation to climate change, since these are characteristics more similar to those existing in southern populations, there are many aspects to consider. First, unexpected effects may occur in different traits. Moreover, the impacts of assisted gene flow will greatly depend on the characteristics of donor and recipient populations, so this strategy will only make sense if the source populations are previously adapted to the environmental conditions now being experienced by the target population (Aitken & Whitlock, 2013; Prieto-Benítez et al., 2021). Ultimately, although the feasibility of these techniques depends on numerous factors and is not an easy question to resolve, our proof of concept using assisted gene flow suggests that these strategies should not be underestimated, as they can be of great use in conserving biodiversity in the current context of climate change. In addition, and considering that this is one of the first studies of its kind, it demonstrates the potential of including genomic analyses to identify the regions being targeted and the real impact on the genome of the populations.

## Acknowledgements

We thank Cristina Poyatos, Pablo Tabarés and Aitor Alameda for the help with the experiments. We also thank Carlos Díaz, José Margalet and Victoria Calvo for the technical support in the CULTIVE facility laboratory greenhouse. This work has been carried out thanks to the financial support of the EVA project (CGL2016-77377-R) of the Spanish Ministry of Science and Innovation. Carlos Lara-Romero was supported by a Juan de la Cierva Incorporación post-doctoral fellowship (Ministerio de Ciencia, Innovación y Universidades: IJC2019-041342-I).

## Notes

### Competing Interest Statement

The authors have declared no competing interest.

