## Supplementary material for "Effects of assisted gene flow on the flowering onset of the annual legume *Lupinus angustifolius* L.: from phenotype to genotype"

Table S1. Populations of *Lupinus angustifolius* L. and common garden site involved in the study. Town, region, geographical coordinates (decimal degrees, WGS84) and climate variables associated to the populations (1985-2015 period) and to the common garden site are shown. May-July period corresponds with the period when the plants are developing fruits and setting seeds. Climate data were obtained from ClimateEU (Marchi et al., 2020).

| **Acronym** | **Town** | **Region** | **Latitude** | **Longitude** | **Elevation (m. a.s.l.)** | **Annual mean temperature (º Celsius)** | **May-July precipitation (mm)** |
|---|---|---|---|---|---|---|---|
| FRO (N) | Zafrón | Central Spain | 41.0241 | -6.0281 | 840 | 12.4 | 92 |
| PIC (N) | Zarapicos | Central Spain | 41.0043 | -5.8130 | 820 | 12.6 | 89 |
| GAR (S) | La Garranchosa | Southern Spain | 38.3257 | -6.4337 | 422 | 16.5 | 64 |
| RIV (S) | Rivera de la Lanchita | Southern Spain | 38.3515 | -6.5760 | 352 | 16.8 | 61 |
| - | Common garden (2017-2020) | Central Spain | 40.3343 | -3.8829 | 690 | 14.9 | 63 |

Table S2. Observed mean ± SD values for the different traits measured and the different lines tested. CFL: control line; GFL: F1 gene flow line; SPL: F2 self-pollination line; BCL: backcross line. SLA: Specific leaflet area; LDMC: leaflet dry matter content.

| | **Flowering onset 2019 (days)** | | **Flowering onset 2020 (days)** | | | **Number of seeds** | | **Seed weigh (mg)** | | **Height (cm)** | | **Biomass (g)** | | **Shoot growth (cm)** | | **SLA ($cm^2/g$)** | | **LDMC (mg/g)** | |
|---|---|---|---|---|---|---|---|---|---|---|---|---|---|---|---|---|---|---|---|
| **Line** | CFL | GFL | CFL | BCL | SPL | CFL | GFL | CFL | GFL | CFL | GFL | CFL | GFL | CFL | GFL | CFL | GFL | CFL | GFL |
| **FRO** | 127,38 ±9,39 | 120,22 ±6,84 | 121,27 ±9,52 | 109,44 ±7,91 | 115,03 ±8,81 | 1239,46 ±679,73 | 1059,49 ±607,55 | 87,95 ±19,59 | 114,97 ±10,75 | 73,57 ±10,75 | 68,72 ±7,98 | 13,81 ±6,85 | 13,43 ±5,91 | 26,39 ±8,64 | 22,84 ±7,85 | 211,69 ±33,22 | 201,03 ±48,78 | 0,13 ±0,01 | 0,13 ±0,02 |
| **PIC** | 128,18 ±9,65 | 120,02 ±6,98 | 119,98 ±13,12 | - | 113,24 ±8,22 | 1003,01 ±481,72 | 954,48 ±405,46 | 80,79 ±23,92 | 97,48 ±17,06 | 74,11 ±11,47 | 72,56 ±9,31 | 13,10 ±5,20 | 14,15 ±4,29 | 28,16 ±10,65 | 23,58 ±8,41 | 215,03 ±29,83 | 201,46 ±46,38 | 0,13 ±0,01 | 0,12 ±0,02 |

Table S3. Effect of the different lines and population on flowering onset, number of seeds, seed weight, height, biomass, shoot growth, SLA and LDMC of *Lupinus angustifolius* plants in the common garden experiment. Estimates and significance level for fixed effects are shown. Genotype was included as random factor. CFL: control line; GFL: F1 gene flow line; SPL: F2 self-pollination line; BCL: backcross line. Missing factors (CFL = Control line and population FRO) are included in the intercept.

| Fixed effects | Parameter value | Standard error | t value | p value | Pseudo-$R^2$ (fixed effects) | Pseudo- $R^2$ (total) |
|---|---|---|---|---|---|---|
| **Flowering onset 2019** | - | - | - | - | 0.100 | 0.122 |
| Intercept | 4.849 | 0.011 | 461.128 | <0.001 | - | - |
| GFL | -0.062 | 0.014 | -4.495 | <0.001 | - | - |
| PIC | 0.003 | 0.013 | 0.251 | 0.802 | - | - |
| **Flowering onset 2020** | - | - | - | - | 0.089 | 0.253 |
| Intercept | 4.798 | 0.015 | 326.774 | <0.001 | - | - |
| BCL | -0.092 | 0.037 | -2.467 | 0.013 | - | - |
| SPL | -0.055 | 0.018 | -3.031 | 0.002 | - | - |
| PIC | -0.013 | 0.018 | -0.714 | 0.475 | - | - |
| **Number of seeds** | - | - | - | - | 0.039 | 0.125 |
| Intercept | 1217.632 | 71.250 | 17.089 | <0.001 | - | - |
| GFL | -107.615 | 91.740 | -1.173 | 0.247 | - | - |
| PIC | -190.473 | 88.460 | -2.153 | 0.036 | - | - |
| **Seed weight** | - | - | - | - | 0.225 | 0.388 |
| Intercept | 89.608 | 2.985 | 30.016 | <0.001 | - | - |
| GFL | 21.602 | 3.947 | 5.743 | <0.001 | - | - |
| PIC | -10.784 | 3.765 | -2.864 | 0.006 | - | - |
| **Height** | - | - | - | - | 0.024 | 0.201 |
| Intercept | 72.992 | 1.390 | 52.494 | <0.001 | - | - |
| GFL | -2.994 | 1.839 | -1.628 | 0.110 | - | - |
| PIC | 1.606 | 1.748 | 0.918 | 0.363 | - | - |
| **Biomass** | - | - | - | - | 0.003 | 0.195 |
| Intercept | 13.538 | 0.823 | 16.448 | <0.001 | - | - |
| GFL | 0.582 | 1.094 | 0.532 | 0.598 | - | - |
| PIC | -0.275 | 1.040 | -0.264 | 0.793 | - | - |
| **Shoot growth** | - | - | - | - | 0.047 | 0.515 |
| Intercept | 26.599 | 1.484 | 17.927 | | - | - |

| | | | | | | |
|---|---|---|---|---|---|---|
| GFL | -4.139 | 2.042 | -2.027 | | - | - |
| PIC | 1.064 | 1.902 | 0.559 | | - | - |
| **SLA** | - | - | - | - | 0.025 | 0.164 |
| Intercept | 212.419 | 4.907 | 43.289 | <0.001 | - | - |
| GFL | -12.522 | 6.537 | -1.915 | 0.061 | - | - |
| PIC | 1.833 | 6.208 | 0.295 | 0.769 | - | - |
| **LDMC** | - | - | - | - | 0.061 | 0.061 |
| Intercept | 0.134 | 0.002 | 79.366 | <0.001 | - | - |
| GFL | -0.001 | 0.002 | -0.496 | 0.622 | - | - |
| PIC | -0.007 | 0.002 | -3.499 | 0.001 | - | - |

Table S4. Chi-square statistic, degrees of freedom and P-values of the Type II Wald chi-square tests of GLMM and LMM analyses to study the effect of selection line, population, and year on flowering onset, number of seeds, seed weight, height, biomass, shoot growth, SLA and LDMC of *Lupinus angustifolius* plants grown in the common garden experiment. Estimates and significance level for fixed effects are shown. Genotype was included as random factor. CFL: control line; GFL: F1 gene flow line; SPL: F2 self-pollination line; BCL: backcross line. Missing factors (CFL = Control line and population FRO) are included in the intercept.

| **Fixed effects** | $X^2$ | **Df** | **Pr (>\|$X^2$\|)** |
|---|---|---|---|
| **Flowering onset 2019** | - | - | - |
| Line | 20.207 | 1 | <0.001 |
| Population | 0.063 | 1 | 0.802 |
| **Flowering onset 2020** | - | - | - |
| Line | 12.998 | 2 | <0.001 |
| Population | 0.510 | 1 | 0.475 |
| **Number of seeds** | - | - | - |
| Line | 1.376 | 1 | 0.241 |
| Population | 4.636 | 1 | 0.031 |
| **Seed weight** | - | - | - |
| Line | 29.959 | 1 | <0.001 |
| Population | 8.204 | 1 | 0.004 |
| **Height** | - | - | - |
| Line | 2.651 | 1 | 0.104 |
| Population | 0.843 | 1 | 0.359 |
| **Biomass** | - | - | - |
| Line | 0.283 | 1 | 0.595 |
| Population | 0.070 | 1 | 0.792 |

| | | | |
|---|---|---|---|
| **Shoot growth** | - | - | - |
| Line | 4.110 | 1 | 0.043 |
| Population | 0.313 | 1 | 0.576 |
| **SLA** | - | - | - |
| Line | 3.669 | 1 | 0.055 |
| Population | 0.0872 | 1 | 0.768 |
| **LDMC** | - | - | - |
| Line | 0.246 | 1 | 0.620 |
| Population | 12.243 | 1 | <0.001 |

Table S5. Posterior mean values, standard errors and 95 % confidence intervals for the different traits and lines of *Lupinus angustifolius* plants grown in a common garden experiment. CFL: control line; GFL: F1 gene flow line; SPL: F2 self-pollination line; BCL: backcross line.

| | **FRO** | | | | **PIC** | | | |
|---|---|---|---|---|---|---|---|---|
| | **Mean** | **Std. error** | **2·5%** | **97·%** | **Mean** | **Std. error** | **2·5%** | **97·%** |
| **Flowering onset 2019** | - | - | - | - | - | - | - | - |
| CFL | 128 | 1.34 | 125 | 130 | 128 | 1.35 | 125 | 131 |
| GFL | 120 | 1.61 | 117 | 123 | 120 | 1.50 | 117 | 123 |
| **Flowering onset 2020** | - | - | - | - | - | - | - | - |
| CFL | 121 | 1.78 | 118 | 125 | 120 | 1.77 | 116 | 123 |
| BCL | 111 | 3.79 | 103 | 118 | 109 | 4.23 | 101 | 118 |
| SPL | 115 | 2.00 | 111 | 119 | 113 | 1.79 | 110 | 117 |
| **Number of seeds** | - | - | - | - | - | - | - | - |
| CFL | 1218 | 71.4 | 1075 | 1360 | 1027 | 70.3 | 887 | 1168 |
| GFL | 1110 | 89.1 | 931 | 1289 | 920 | 82.2 | 754 | 1085 |
| **Seed weight** | - | - | - | - | - | - | - | - |
| CFL | 89.60 | 2.99 | 83.60 | 95.60 | 78.80 | 2.99 | 72.80 | 84.80 |
| GFL | 111.20 | 3.88 | 103.40 | 119.00 | 100.40 | 3.54 | 93.30 | 107.50 |
| **Height** | - | - | - | - | - | - | - | - |
| CFL | 73.00 | 1.39 | 70.20 | 75.80 | 74.60 | 1.38 | 71.80 | 77.40 |
| GFL | 70.00 | 1.79 | 66.40 | 73.60 | 71.60 | 1.67 | 68.30 | 75.00 |
| **Biomass** | - | - | - | - | - | - | - | - |
| CFL | 13.50 | 0.83 | 11.90 | 15.20 | 13.30 | 0.83 | 11.60 | 14.90 |
| GFL | 14.40 | 1.07 | 12.00 | 16.30 | 13.80 | 0.99 | 11.90 | 15.80 |
| **Shoot growth** | - | - | - | - | - | - | - | - |
| CFL | 26.60 | 1.48 | 23.60 | 29.60 | 27.70 | 1.50 | 24.70 | 30.70 |

| | | | | | | | | |
|---|---|---|---|---|---|---|---|---|
| GFL | 22.50 | 2.01 | 18.40 | 26.50 | 23.50 | 1.87 | 19.80 | 27.30 |
| **SLA** | - | - | - | - | - | - | - | - |
| CFL | 212 | 4.91 | 203 | 222 | 214 | 4.93 | 204 | 224 |
| GFL | 200 | 6.36 | 187 | 213 | 202 | 5.94 | 190 | 214 |
| **LDMC** | - | - | - | - | - | - | - | - |
| CFL | 0.13 | 0.00 | 0.13 | 0.14 | 0.13 | 0.00 | 0.12 | 0.13 |
| GFL | 0.13 | 0.00 | 0.13 | 0.14 | 0.13 | 0.00 | 0.12 | 0.13 |

Table S6. Minimum allele frequency (MAF), False Discovery Rate (FDR), $F_{ST}$ statistic and $F_{ST}$-FDR values for each SNP identified as outlier in the genomic analyses.

| | | | | | | | CFL | GFL |
|---|---|---|---|---|---|---|---|---|
| **Population** | **SNP** | **Gene** | **Protein** | **FDR** | **Fst** | **Fst-FDR** | **MAF** | **MAF** |
| FRO | NC_032009.1_17378625 | LOC109350320 | XP_019447099.1 | 0.010 | 0.313 | <0.001 | 0.067 | 0.467 |
| FRO | NC_032009.1_19045875 | LOC109351227 | XP_019448173.1 | 0.006 | 0.348 | <0.001 | 0.067 | 0.500 |
| FRO | NC_032009.1_19129990 | LOC109351285 | XP_019448262.1 | 0.006 | 0.348 | <0.001 | 0.067 | 0.500 |
| FRO | NC_032009.1_3656339 | LOC109347866 | XP_019443510.1 | 0.010 | 0.313 | <0.001 | 0.067 | 0.467 |
| FRO | NC_032009.1_3656428 | LOC109347866 | XP_019443510.1 | 0.010 | 0.313 | <0.001 | 0.067 | 0.467 |
| FRO | NC_032010.1_1906478 | LOC109331801 | XP_019422070.1 | 0.023 | 0.266 | 0.001 | 0.100 | 0.467 |
| FRO | NC_032010.1_4793322 | LOC109328838 | XP_019417994.1 | 0.045 | 0.212 | 0.003 | 0.133 | 0.464 |
| FRO | NC_032010.1_4793895 | LOC109328838 | XP_019417994.1 | 0.048 | 0.214 | 0.003 | 0.133 | 0.467 |
| FRO | NC_032011.1_9190057 | LOC109343327 | XP_019437115.1 | 0.033 | 0.247 | 0.001 | 0.033 | 0.333 |
| FRO | NC_032011.1_9190442 | LOC109343327 | XP_019437115.1 | 0.033 | 0.247 | 0.001 | 0.033 | 0.333 |
| FRO | NC_032012.1_1017139 | LOC109345004 | XP_019439298.1 | 0.009 | 0.309 | <0.001 | 0.250 | 0.700 |
| FRO | NC_032012.1_2222950 | LOC109345069 | XP_019439392.1 | 0.001 | 0.489 | <0.001 | 0.067 | 0.607 |
| FRO | NC_032012.1_946558 | LOC109344999 | XP_019439290.1 | 0.002 | 0.410 | <0.001 | 0.033 | 0.500 |
| FRO | NC_032013.1_1922165 | LOC109347401 | XP_019442778.1 | 0.023 | 0.271 | 0.001 | 0.100 | 0.467 |
| FRO | NC_032013.1_1930095 | LOC109347401 | XP_019442778.1 | 0.023 | 0.271 | 0.001 | 0.100 | 0.467 |
| FRO | NC_032013.1_1939091 | LOC109347400 | XP_019442777.1 | 0.048 | 0.219 | 0.002 | 0.133 | 0.467 |
| FRO | NC_032013.1_1941466 | LOC109347400 | XP_019442777.1 | 0.048 | 0.219 | 0.002 | 0.133 | 0.467 |
| FRO | NC_032013.1_1941529 | LOC109347400 | XP_019442777.1 | 0.048 | 0.219 | 0.002 | 0.133 | 0.467 |
| FRO | NC_032013.1_1943836 | LOC109347400 | XP_019442777.1 | 0.023 | 0.271 | 0.001 | 0.100 | 0.467 |
| FRO | NC_032013.1_22007083 | LOC109348120 | XP_019443898.1 | 0.029 | 0.238 | 0.001 | 0.033 | 0.321 |
| FRO | NC_032013.1_22008378 | LOC109348120 | XP_019443898.1 | 0.033 | 0.240 | 0.001 | 0.033 | 0.333 |
| FRO | NC_032013.1_22020240 | LOC109348121 | XP_019443899.1 | 0.033 | 0.247 | 0.001 | 0.033 | 0.333 |
| FRO | NC_032014.1_29566731 | LOC109350790 | XP_019447641.1 | 0.035 | 0.229 | 0.002 | 0.467 | 0.833 |
| FRO | NC_032019.1_35464640 | LOC109360702 | XP_019461304.1 | 0.048 | 0.209 | 0.003 | 0.133 | 0.467 |
| FRO | NC_032021.1_13472971 | LOC109325863 | XP_019414003.1 | 0.040 | 0.227 | 0.002 | 0.433 | 0.100 |
| FRO | NC_032024.1_1858436 | LOC109329933 | XP_019419387.1 | 0.008 | 0.326 | <0.001 | 0.100 | 0.533 |
| FRO | NC_032025.1_19286248 | LOC109331033 | XP_019420861.1 | 0.002 | 0.457 | <0.001 | 0.133 | 0.679 |
| FRO | NC_032025.1_19287622 | LOC109331033 | XP_019420861.1 | 0.035 | 0.243 | 0.001 | 0.167 | 0.533 |

| | | | | | | | | |
|---|---|---|---|---|---|---|---|---|
| FRO | NC_032028.1_459890 | LOC109335596 | XP_019427286.1 | 0.01 | 0.323 | <0.001 | 0.067 | 0.467 |
| FRO | NC_032028.1_459979 | LOC109335596 | XP_019427286.1 | 0.023 | 0.266 | 0.001 | 0.100 | 0.467 |
| FRO | NC_032028.1_460030 | LOC109335596 | XP_019427286.1 | 0.006 | 0.357 | <0.001 | 0.067 | 0.500 |
| FRO | NC_032028.1_460163 | LOC109335596 | XP_019427286.1 | 0.021 | 0.265 | 0.001 | 0.100 | 0.464 |
| FRO | NC_032028.1_460389 | LOC109335596 | XP_019427286.1 | 0.048 | 0.204 | 0.003 | 0.133 | 0.467 |
| FRO | NC_032028.1_460639 | LOC109335596 | XP_019427286.1 | 0.003 | 0.392 | <0.001 | 0.067 | 0.533 |
| FRO | NW_017722081.1_3838 | LOC109338905 | XP_019431800.1 | 0.023 | 0.276 | 0.001 | 0.100 | 0.467 |
| FRO | NW_017722081.1_4164 | LOC109338905 | XP_019431800.1 | 0.013 | 0.314 | <0.001 | 0.100 | 0.500 |
| PIC | NC_032009.1_17378625 | LOC109350320 | XP_019447099.1 | 0.045 | 0.315 | 0.001 | 0.033 | 0.400 |
| PIC | NC_032009.1_19045875 | LOC109351227 | XP_019448173.1 | 0.045 | 0.315 | 0.001 | 0.033 | 0.400 |
| PIC | NC_032009.1_19129990 | LOC109351285 | XP_019448262.1 | 0.045 | 0.315 | 0.001 | 0.033 | 0.400 |
| PIC | NC_032009.1_3656339 | LOC109347866 | XP_019443510.1 | 0.040 | 0.328 | 0.001 | 0.067 | 0.467 |
| PIC | NC_032009.1_3656428 | LOC109347866 | XP_019443510.1 | 0.040 | 0.328 | 0.001 | 0.067 | 0.467 |
| PIC | NC_032010.1_1906478 | LOC109331801 | XP_019422070.1 | 0.029 | 0.358 | <0.001 | 0.033 | 0.433 |
| PIC | NC_032010.1_4793322 | LOC109328838 | XP_019417994.1 | 0.029 | 0.358 | <0.001 | 0.033 | 0.433 |
| PIC | NC_032010.1_4793895 | LOC109328838 | XP_019417994.1 | 0.029 | 0.358 | <0.001 | 0.033 | 0.433 |
| PIC | NC_032011.1_9190057 | LOC109343327 | XP_019437115.1 | 0.040 | 0.321 | 0.001 | <0.001 | 0.333 |
| PIC | NC_032011.1_9190442 | LOC109343327 | XP_019437115.1 | 0.040 | 0.321 | 0.001 | <0.001 | 0.333 |
| PIC | NC_032012.1_1017139 | LOC109345004 | XP_019439298.1 | 0.040 | 0.315 | 0.001 | 0.433 | 0.867 |
| PIC | NC_032012.1_2222950 | LOC109345069 | XP_019439392.1 | 0.032 | 0.349 | <0.001 | 0.367 | 0.833 |
| PIC | NC_032012.1_946558 | LOC109344999 | XP_019439290.1 | 0.029 | 0.358 | <0.001 | 0.033 | 0.433 |
| PIC | NC_032013.1_1922165 | LOC109347401 | XP_019442778.1 | 0.040 | 0.328 | 0.001 | 0.067 | 0.467 |
| PIC | NC_032013.1_1930095 | LOC109347401 | XP_019442778.1 | 0.040 | 0.328 | 0.001 | 0.067 | 0.467 |
| PIC | NC_032013.1_1939091 | LOC109347400 | XP_019442777.1 | 0.040 | 0.328 | 0.001 | 0.067 | 0.467 |
| PIC | NC_032013.1_1941466 | LOC109347400 | XP_019442777.1 | 0.040 | 0.328 | 0.001 | 0.067 | 0.467 |
| PIC | NC_032013.1_1941529 | LOC109347400 | XP_019442777.1 | 0.040 | 0.328 | 0.001 | 0.067 | 0.467 |
| PIC | NC_032013.1_1943836 | LOC109347400 | XP_019442777.1 | 0.040 | 0.328 | 0.001 | 0.067 | 0.467 |
| PIC | NC_032013.1_22007083 | LOC109348120 | XP_019443898.1 | 0.040 | 0.321 | 0.001 | <0.001 | 0.333 |
| PIC | NC_032013.1_22008378 | LOC109348120 | XP_019443898.1 | 0.040 | 0.314 | 0.001 | <0.001 | 0.333 |
| PIC | NC_032013.1_22020240 | LOC109348121 | XP_019443899.1 | 0.040 | 0.321 | 0.001 | <0.001 | 0.333 |
| PIC | NC_032014.1_29566731 | LOC109350790 | XP_019447641.1 | 0.040 | 0.300 | 0.001 | 0.667 | 1 |
| PIC | NC_032019.1_35464640 | LOC109360702 | XP_019461304.1 | 0.040 | 0.328 | 0.001 | 0.067 | 0.467 |
| PIC | NC_032021.1_13472971 | LOC109325863 | XP_019414003.1 | 0.029 | 0.385 | <0.001 | 0.900 | 0.433 |

| | | | | | | | | |
|---|---|---|---|---|---|---|---|---|
| PIC | NC_032024.1_1858436 | LOC109329933 | XP_019419387.1 | 0.040 | 0.318 | 0.001 | 0.067 | 0.467 |
| PIC | NC_032025.1_19286248 | LOC109331033 | XP_019420861.1 | 0.040 | 0.324 | 0.001 | 0.107 | 0.533 |
| PIC | NC_032025.1_19287622 | LOC109331033 | XP_019420861.1 | 0.040 | 0.328 | 0.001 | 0.067 | 0.467 |
| PIC | NC_032028.1_459890 | LOC109335596 | XP_019427286.1 | 0.009 | 0.429 | <0.001 | <0.001 | 0.433 |
| PIC | NC_032028.1_459979 | LOC109335596 | XP_019427286.1 | 0.009 | 0.459 | <0.001 | <0.001 | 0.467 |
| PIC | NC_032028.1_460030 | LOC109335596 | XP_019427286.1 | 0.009 | 0.429 | <0.001 | <0.001 | 0.433 |
| PIC | NC_032028.1_460163 | LOC109335596 | XP_019427286.1 | 0.009 | 0.423 | <0.001 | <0.001 | 0.433 |
| PIC | NC_032028.1_460389 | LOC109335596 | XP_019427286.1 | 0.005 | 0.555 | <0.001 | <0.001 | 0.567 |
| PIC | NC_032028.1_460639 | LOC109335596 | XP_019427286.1 | 0.021 | 0.393 | <0.001 | <0.001 | 0.400 |
| PIC | NW_017722081.1_3838 | LOC109338905 | XP_019431800.1 | 0.029 | 0.358 | <0.001 | 0.033 | 0.433 |
| PIC | NW_017722081.1_4164 | LOC109338905 | XP_019431800.1 | 0.029 | 0.358 | <0.001 | 0.033 | 0.433 |

Table S7. Functional annotation of the 36 SNPs identified as outliers in the genomic analyses.

| Name | Protein name | Protein ID | GO biological process |
|---|---|---|---|
| NC_032009.1_17378625 | axial regulator YABBY 1-like | XP_019447099.1 | Flower development (flowering) |
| NC_032009.1_19045875 | uncharacterized protein | XP_019448173.1 | Meiotic nuclear division (reproduction) |
| NC_032009.1_19129990 | xyloglucan endotransglucosylase/hydrolase protein 28 | XP_019448262 | Stamen filament development (flowering) |
| NC_032009.1_3656339 | 3-oxoacyl-[acyl-carrier-protein] synthase II, chloroplastic-like | XP_019443510 | Response to cold (response to abiotic stress) |
| NC_032009.1_3656428 | 3-oxoacyl-[acyl-carrier-protein] synthase II, chloroplastic-like | XP_019443510.1 | Response to cold (response to abiotic stress) |
| NC_032010.1_1906478 | nitrogen regulatory protein P-II homolog | XP_019422070.1 | Regulation of nitrogen utilization (nitrogen) |
| NC_032010.1_4793322 | rop guanine nucleotide exchange factor 12-like | XP_019417994.1 | Pollen tube growth (flowering) |
| NC_032010.1_4793895 | rop guanine nucleotide exchange factor 12-like | XP_019417994.1 | Pollen tube growth (flowering) |
| NC_032011.1_9190057 | chaperone protein dnaJ GFA2, mitochondrial-like isoform X3 | XP_019437115.1 | Pollination (reproduction) |
| NC_032011.1_9190442 | chaperone protein dnaJ GFA2, mitochondrial-like isoform X3 | XP_019437115.1 | Pollination (reproduction) |
| NC_032012.1_1017139 | isoleucine--tRNA ligase, chloroplastic/mitochondrial | XP_019439298.1 | Ovule development (flowering) |
| NC_032012.1_2222950 | cytochrome P450 90A1-like | XP_019439392.1 | Anther differentiation (flowering) |
| NC_032012.1_946558 | protein pleiotropic regulatory locus 1-like | XP_019439290.1 | Cotyledon development (reproduction) |

| | | | |
|---|---|---|---|
| NC_032013.1_1922165 | cell division cycle protein 27 homolog B-like isoform X1 | XP_019442778.1 | Root meristem specification (growth) |
| NC_032013.1_1930095 | cell division cycle protein 27 homolog B-like isoform X1 | XP_019442778.1 | Root meristem specification (growth) |
| NC_032013.1_1939091 | cell division cycle protein 27 homolog B-like | XP_019442777.1 | Root meristem specification (growth) |
| NC_032013.1_1941466 | cell division cycle protein 27 homolog B-like | XP_019442777.1 | Root meristem specification (growth) |
| NC_032013.1_1941529 | cell division cycle protein 27 homolog B-like | XP_019442777.1 | Root meristem specification (growth) |
| NC_032013.1_1943836 | cell division cycle protein 27 homolog B-like | XP_019442777.1 | Root meristem specification (growth) |
| NC_032013.1_22007083 | chromatin remodeling protein EBS-like | XP_019443898.1 | Regulation of long-day photoperiodism (flowering) |
| NC_032013.1_22008378 | chromatin remodeling protein EBS-like | XP_019443898.1 | Regulation of long-day photoperiodism (flowering) |
| NC_032013.1_22020240 | chromatin remodeling protein EBS-like | XP_019443899.1 | Regulation of long-day photoperiodism (flowering) |
| NC_032014.1_29566731 | Cyclic nucleotide-binding domain-containing protein; putative cyclic nucleotide-gated ion channel 8 isoform X2 | OIW09331.1; XP_019447642.1 | Pollen tube growth (flowering) |
| NC_032019.1_35464640 | ubiquitin protein ligase | OIW01082.1 | Chromatin organization |
| NC_032021.1_13472971 | armadillo repeat-containing protein LFR | XP_019414003.1 | Anther development (flowering) |
| NC_032024.1_1858436 | nuclear export mediator factor NEMF | XP_019419388.1 | Cold acclimation (response to abiotic stress) |
| NC_032025.1_19286248 | protein FLOWERING LOCUS T-like | OIV93971 1; XP_019420861 | Response to short-day photoperiodism (flowering) |

| | | | |
|---|---|---|---|
| NC_032025.1_19287622 | protein FLOWERING LOCUS T-like | XP_019420861.1 | Response to short-day photoperiodism (flowering) |
| NC_032028.1_459890 | transcription factor RF2b-like | OIV91432.1; XP_019427286.1 | Response to sulfate (response to abiotic stress) |
| NC_032028.1_459979 | transcription factor RF2b-like | OIV91432.1; XP_019427286.2 | Response to sulfate (response to abiotic stress) |
| NC_032028.1_460030 | transcription factor RF2b-like | OIV91432.1; XP_019427286.3 | Response to sulfate (response to abiotic stress) |
| NC_032028.1_460163 | transcription factor RF2b-like | OIV91432.1; XP_019427286.4 | Response to sulfate (response to abiotic stress) |
| NC_032028.1_460389 | transcription factor RF2b-like | OIV91432.1; XP_019427286.5 | Response to sulfate (response to abiotic stress) |
| NC_032028.1_460639 | transcription factor RF2b-like | XP_019427286.1 | Response to sulfate (response to abiotic stress) |
| NW_017722081.1_3838 | ER lumen protein-retaining receptor | XP_019431801.1_1 | Meiotic nuclear division (reproduction) |
| NW_017722081.1_4164 | ER lumen protein-retaining receptor | XP_019431801.1_1 | Meiotic nuclear division (reproduction) |

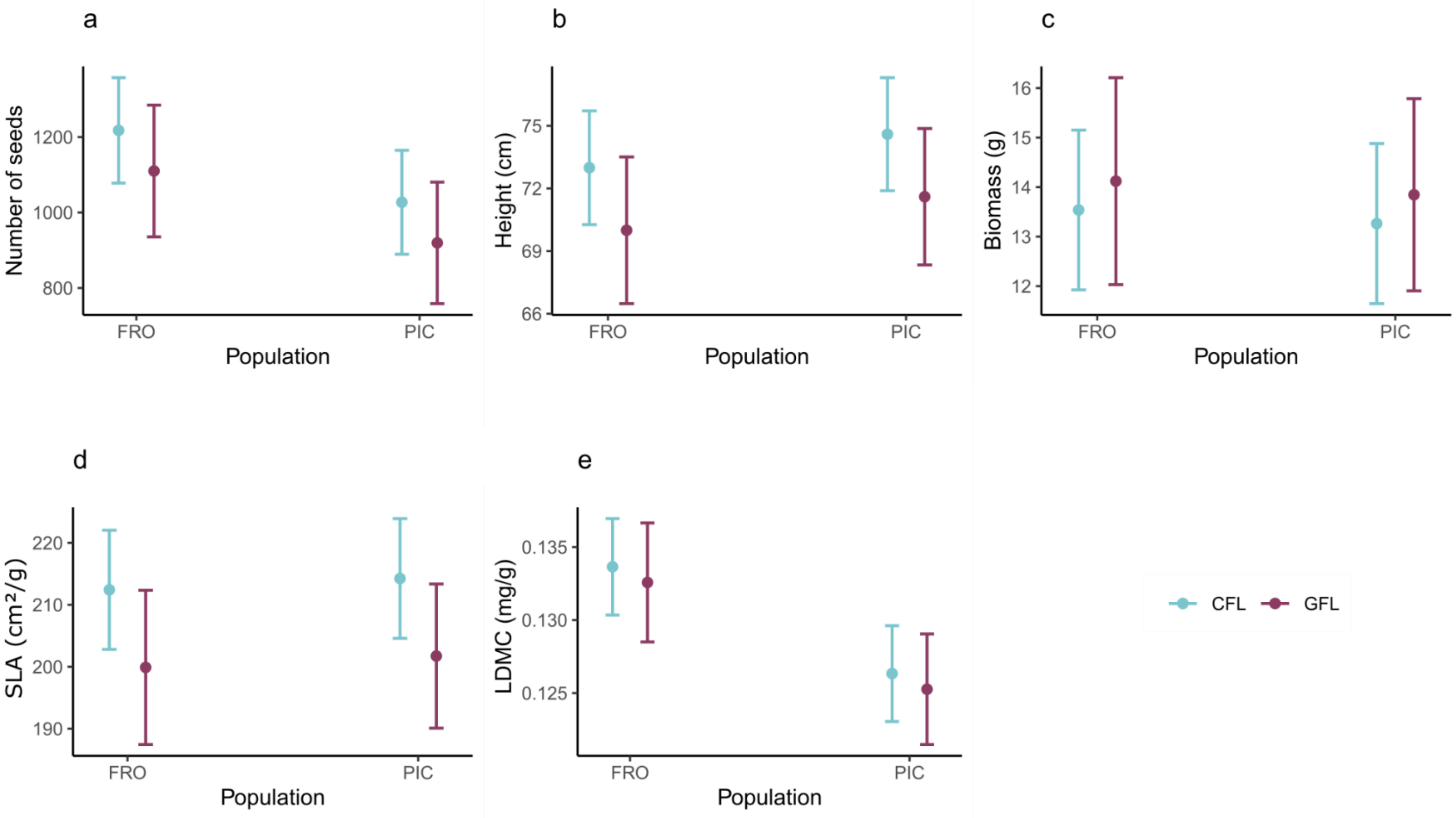


Figure S1. Effect of the gene flow line (GFL) on the different traits of *Lupinus angustifolius* L.: a) number of seeds, b) height, c) biomass, d) SLA, e) LDMC. Dots and bars represent the predicted mean from the LMM model with a Gaussian distribution and the 95 % confidence intervals. Differences between the gene flow line and the control line were non-significant.

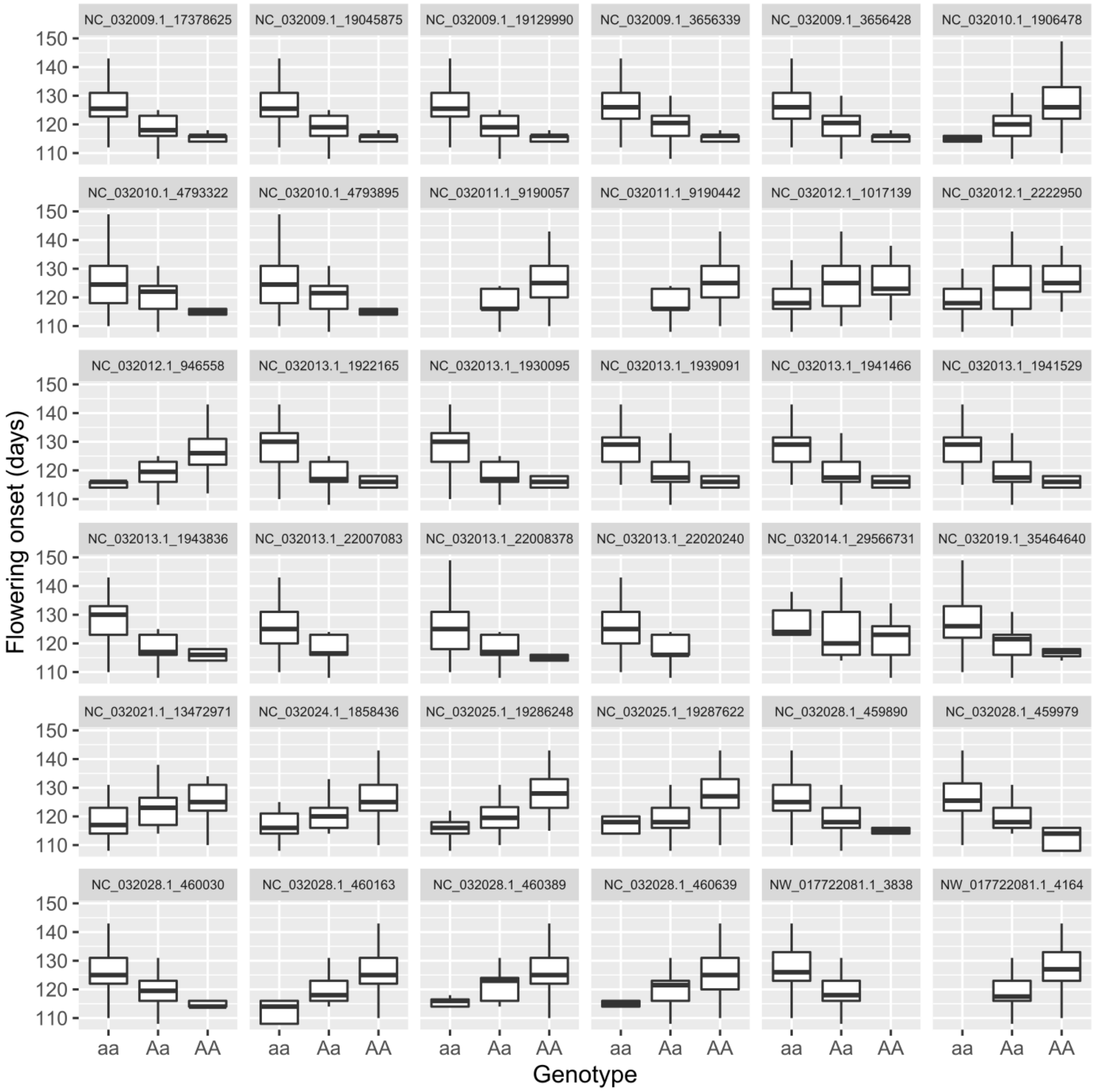


Figure S2. Distribution of flowering onset (days) according to the genotypes (homozygous dominant, recessive and heterozygous) for the 36 significant SNPs detected.

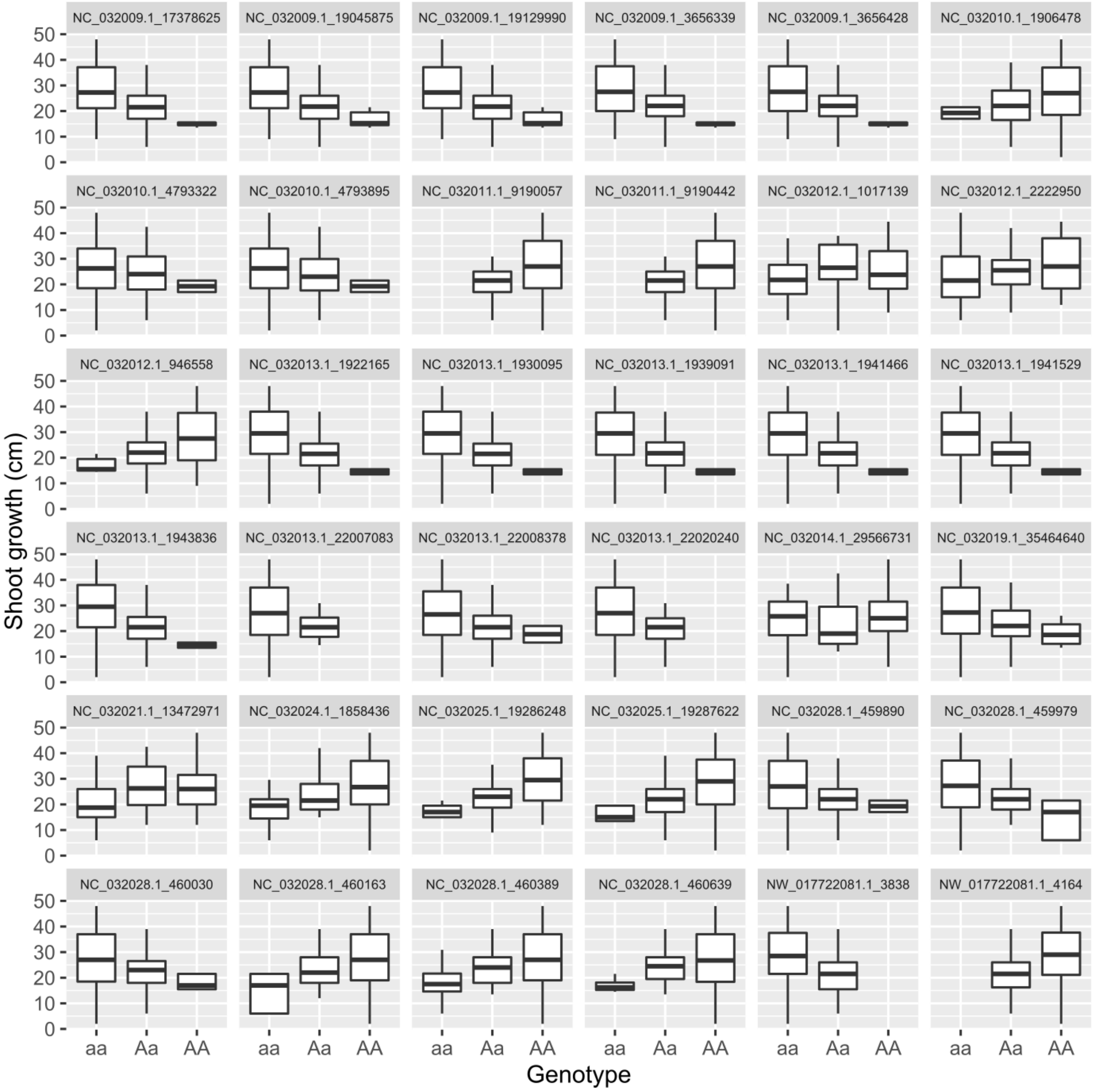


Figure S3. Distribution of the shoot growth (cm) according to the genotypes (homozygous dominant, recessive and heterozygous) for the 36 significant SNPs detected.

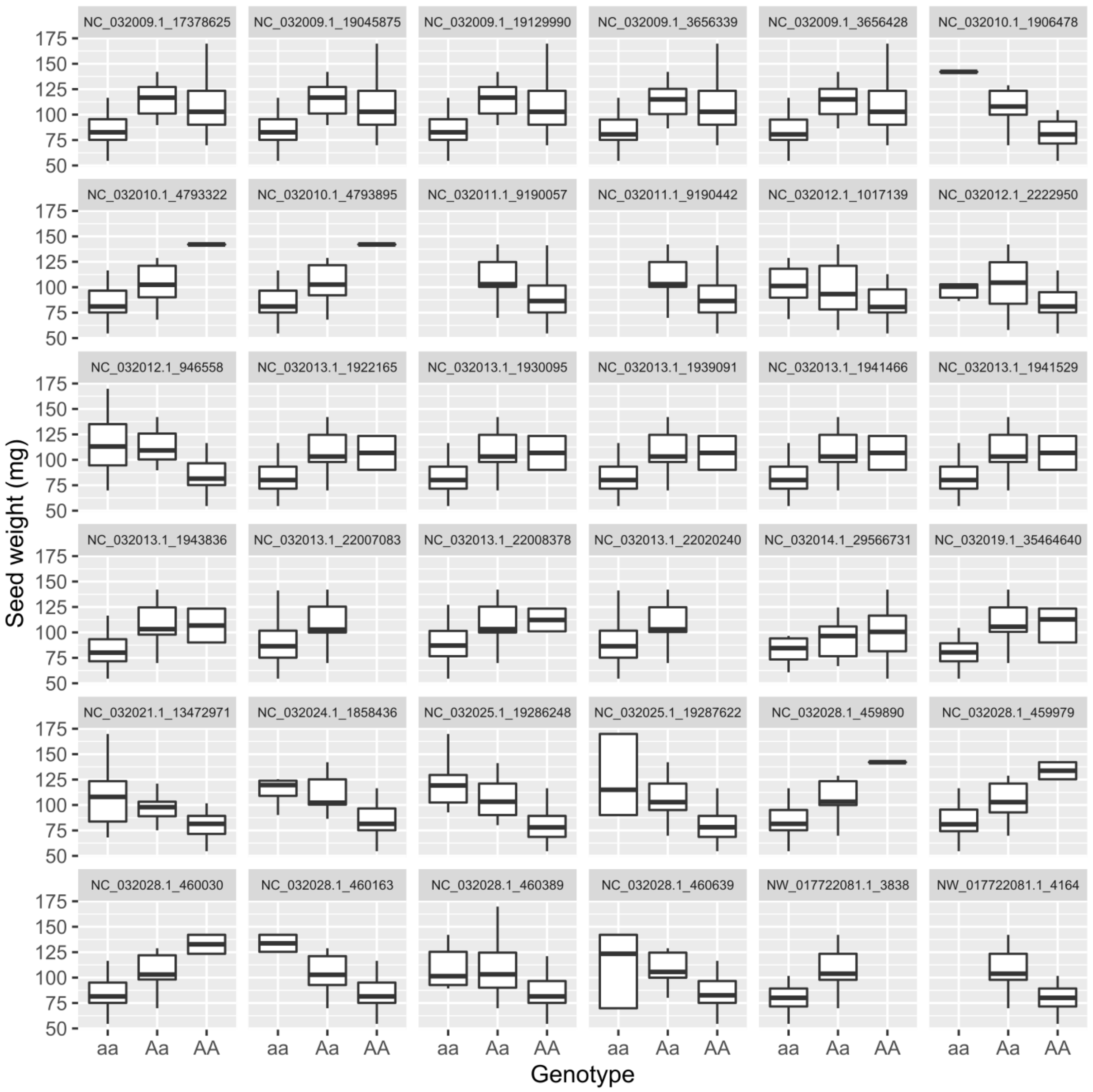


Figure S4. Distribution of the seed weight (mg) according to the genotypes (homozygous dominant, recessive and heterozygous) for the 36 significant SNPs detected.

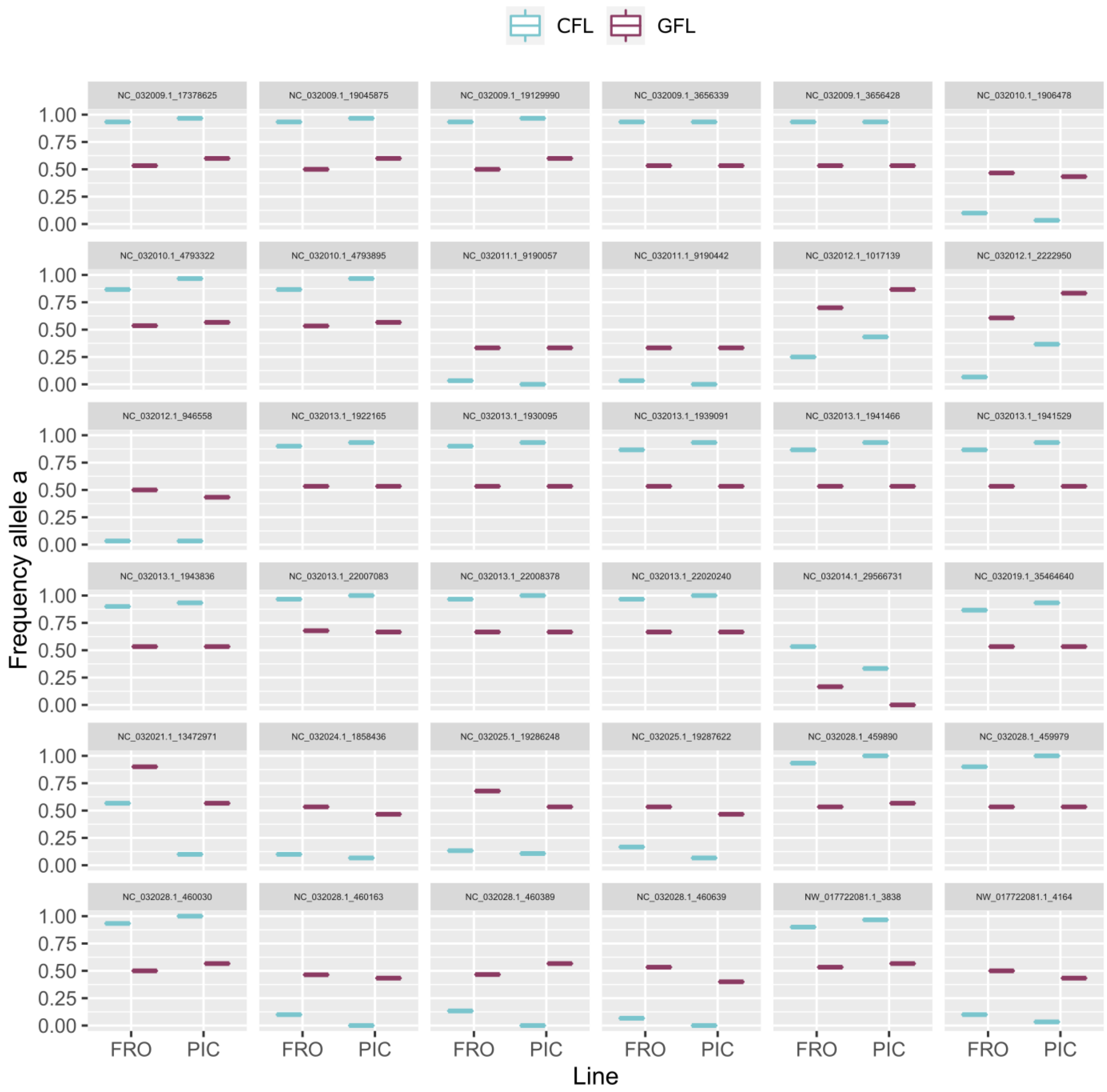


Figure S5. Changes in allele frequencies between the control treatment (blue) and the gene flow treatment (purple) for both populations (FRO and PIC) for the 36 significant SNPs detected.
